# Asymmetric coding of reward prediction errors in human insula and dorsomedial prefrontal cortex

**DOI:** 10.1101/2022.12.07.519496

**Authors:** Colin W. Hoy, David R. Quiroga-Martinez, David King-Stephens, Kenneth D. Laxer, Peter Weber, Jack J. Lin, Robert T. Knight

**Affiliations:** Department of Neurology, University of California, San Francisco; Helen Wills Neuroscience Institute, University of California, Berkeley; Center for Music in the Brain, Aarhus University & The Royal Academy of Music, Aarhus, Denmark; Department of Neurology and Neurosurgery, California Pacific Medical Center, San Francisco, California; Department of Neurology, Yale School of Medicine; Department of Neurology, University of California, Davis; Center for Mind and Brain, University of California, Davis; Department of Psychology, University of California, Berkeley

## Abstract

The signed value and unsigned salience of reward prediction errors (RPEs) are critical to understanding reinforcement learning (RL) and cognitive control. Dorsomedial prefrontal cortex (dMPFC) and insula (INS) are key regions for integrating reward and surprise information, but conflicting evidence for both signed and unsigned activity has led to competing proposals for the nature of RPE representations in these brain areas. Recently, the distributional RL theory (dRL) has been used to explain RPE coding diversity in the rodent midbrain by proposing that dopaminergic neurons have differential sensitivity to positive and negative RPEs. Here, we use intracranially recorded high frequency activity (HFA) to show that this ***asymmetric scaling*** strategy captures RPE coding diversity in human dMPFC and INS. We found neural populations responding to valence-specific positive and negative RPEs, as well as unsigned RPE salience, which are spatially interleaved within each region. Furthermore, directional connectivity estimates suggest a leading role of INS in communicating positive and unsigned RPEs to dMPFC. These findings support asymmetric scaling across distinct but intermingled neural populations as a core principle in RPE coding, expand the scope of dRL, and reconcile longstanding theoretical debates on the role of dMPFC and INS in RL and cognitive control.

## INTRODUCTION

Adaptive behavior requires predicting the stimuli or actions associated with valuable outcomes. Surprising violations of these predictions (i.e., reward prediction errors, or RPEs) are used to learn and update such associations^1^. The scalar value of RPEs has both signed valence (better or worse than expected?), which reinforces either approach or avoidance behavior, as well as unsigned salience (absolute magnitude, or total surprise) that drives motivation, arousal, and motor preparation^2^. Dorsomedial prefrontal cortex (dMPFC) and the insula (INS) are two key brain regions that respond to both RPE valence and salience^3,4^. These areas have strong anatomical and functional connections and together form the salience network (also referred to as cingulo-opercular control network), which is involved in performance monitoring and integrating feedback to adjust cognitive control^5–9^. However, conflicting reports linking dMPFC and INS activity to a diverse range of signed and unsigned RPE signals have fueled long standing theoretical debates about their role in reward learning and cognitive control.

Theories of salience network function have focused primarily on explaining dMPFC activity and can be classified into three families. In one family, theories posit dMPFC encodes positive and negative RPEs together as a “common currency” value or utility signal to inform action selection^10–13^. A second family suggests dMPFC is specialized for processing negative RPEs to coordinate responses to threats and pain^14–16^. A family of alternative theories argue that dMPFC primarily responds to various unsigned salience signals, either to adjust cognitive control^17,18^, orient towards novel or surprising stimuli^19^, or track uncertainty in the environment related to exploration and foraging^20^.

One barrier to addressing these competing theories is that many studies assume positive and negative RPEs are represented together on a symmetric, linear scale relative to a single mean expected value. This classical reinforcement learning (RL) model is partly inspired by foundational observations of dopaminergic neurons that increase their firing rate to positive RPEs and decrease it to negative RPEs^21,22^. However, recent single unit studies in animals have demonstrated that different subpopulations of midbrain dopaminergic neurons separately code for positive RPEs, negative RPEs, and unsigned RPE salience^23–27^. Therefore, alternative computational models are required to account for the unexplained RPE coding diversity in dopaminergic neurons and reconcile theoretical debates on dMPFC and salience network function.

The distributional RL theory (dRL) has offered an explanation for the diversity in neural RPE signals in subcortical dopaminergic neurons in rodents^28^. dRL posits each neuron represents a different expected value with varying levels of optimism and pessimism, such that the full distribution of possible outcomes is encoded by the population. This is achieved by allowing neurons to have valence-specific learning rates, so that they are differentially sensitive to positive and negative reward outcomes. This feature, known as ***asymmetric scaling***, captures the heterogeneity of reward and punishment signals in subcortical dopaminergic neurons in rodents and has improved performance of deep RL models^28–30^. However, whether asymmetric scaling underlies signed and unsigned RPE coding in the human cortex is unclear.

Another challenge for assessing theories of neural RPE coding is that non-human primate studies indicate populations of single units tracking positive and negative RPEs are intermingled within dMPFC^11,31–33^. These populations can represent information with heterogeneous coding schemes using both increases and decreases of activity^34,35^, which can confound valence-specific responses. Common analysis strategies in human cognitive neuroscience average activity within a region and are thus not well suited for resolving overlapping circuits with opposite valence coding and/or directionality of activity changes, particularly for data with lower spatial resolution such as scalp electroencephalography (EEG).

Intracranial EEG (iEEG) recordings with high spatiotemporal resolution overcome some of the limitations of non-invasive human methods. A recent human iEEG study reported an anatomical dissociation between positive and negative RPE processing across regions associated with value-based decision making, including a bias for negative RPEs in anterior INS^36^. However, this study did not record from dMPFC and focused on region-level analyses that may obscure the different contributions of overlapping circuits within each region.

A final important challenge in elucidating dMPFC function is to understand the flow of information in the salience network. Traditionally, dMPFC has been regarded as a control hub where information about task performance, conflict and reward is computed. However, similar representations of signed and unsigned RPE variables are also reported in the relatively less studied INS^37–41^, and recent evidence suggests that the INS may lead information transfer to dMPFC^42–44^. Experimental designs and computational models that dissociate signed (positive and/or negative) and unsigned RPEs are required to elucidate the role of the INS in RPE processing and communication.

Here, we bridge these gaps between species, recording modalities, analysis methods, and computational models by testing whether the asymmetric scaling principle inspired by dRL can dissociate signed positive and negative, as well as unsigned RPE responses in local populations of human dMPFC and INS. We recorded iEEG data from 10 epilepsy patients with combined coverage in dMPFC and INS while they performed a target time task that used difficulty to manipulate expected outcomes and provide the critical dissociation of RPE valence and salience^45–47^. Using high-frequency activity (HFA) power as a marker of local population dynamics^48–51^, we compared the performance of three different linear mixed models in explaining single-trial dMPFC and INS responses to positive, negative, and neutral feedback during the task. We contrasted a ***signed*** model with linear RPE *value* as a classical RL predictor; an ***unsigned*** model with RPE *salience* (i.e., absolute RPE magnitude) as a surprise-related predictor; and an ***asymmetric*** model in which absolute negative and positive RPE magnitude were entered as separate predictors. In the asymmetric model, different regression slopes for positive and negative predictors would indicate asymmetric scaling of RPEs.

We found that the asymmetric model explained RPE signals in dMPFC and INS better than traditional signed and unsigned models. Furthermore, individual electrode sites showed differential responsiveness to positive and negative RPEs, such that spatially intermingled neuronal populations separately encoded positive RPEs, negative RPEs, and unsigned RPE salience. Signed RPE value coding was relatively rare, arguing against theories claiming dMPFC primarily represents RPEs in a symmetric, linear scheme. Moreover, in disagreement with classical RL and dRL, a substantial portion of channel sites increased activity to negative RPE and decreased it to positive RPE. This “inverted” coding scheme invites future work to expand the scope of dRL. Finally, directed connectivity measures suggested positive and unsigned RPE information was primarily transmitted from INS to dMPFC, while negative and signed RPEs showed limited connectivity modulations. These results resolve competing theories of dMPFC function by demonstrating that asymmetric scaling enables both valence-specific and unsigned RPE salience signals that coexist within overlapping dMPFC and INS circuits, while also suggesting that INS plays a leading role in positive and unsigned RPE processing within the salience network.

## RESULTS

We collected behavioral data from 10 patients while recording from implanted SEEG and ECoG electrodes in dMPFC (primarily mid-cingulate cortex with some supplementary motor complex and anterior cingulate sites), and INS (Fig. 1a; see Table 1 for patient demographics, electrode coverage, and behavior). These patients performed a Target Time task that dissociates valenced RPE value and non-valenced RPE magnitude by using task difficulty manipulations to modulate reward expectations in an interval timing paradigm (Fig. 1b). Error tolerance was adjusted after each trial by two staircase algorithms to clamp accuracy at 74.4 ± 6.9% and 19.5 ± 2.6% (mean ± SD) in easy and hard blocks, respectively. This design dissociates outcome valence and probability by manipulating whether wins or losses are surprising, allowing separation of valenced and non-valenced RPE features. Four patients performed a version of the task that delivered neutral outcomes with no RT feedback on 12% of trials as an additional source of surprise.

**Table 1:** Patient demographics, electrode coverage, and behavior. For Button colum, “Kb” indicates responses were collected using the space bar on the built-in laptop keyboard, while “RTBox” indicates a USB button box was used. *For IR87, three runs used the RTbox device, while the keyboard was used to capture responses on the fourth run.

| SBJ | Age (years) | Sex | Task Version | Button | Number of Electrodes |  | Number of Trials |  | Accuracy (%) |  |
| --- | --- | --- | --- | --- | --- | --- | --- | --- | --- | --- |
|  |  |  |  |  | dMPF C | Insula | Easy | Hard | Easy | Hard |
| S01 | 24 | M | 1.8.7 | Kb | 16 | 0 | 299 | 297 | 81.6 | 23.9 |
| S02 | 23 | M | 1.8.2 | Kb | 16 | 13 | 140 | 138 | 67.9 | 18.8 |
| S03 | 27 | M | 1.8.7 | Kb | 18 | 3 | 132 | 149 | 75.0 | 20.1 |
| S04 | 28 | M | 1.8.7 | Kb | 17 | 6 | 145 | 149 | 62.8 | 21.5 |
| S05 | 57 | M | 1.8.7 | Kb | 5 | 5 | 147 | 147 | 70.1 | 19.7 |
| S06 | 47 | M | 1.8.8 | Kb | 8 | 6 | 141 | 133 | 77.3 | 21.8 |
| S07 | 21 | F | 2.4.5 | RTBox | 9 | 8 | 144 | 143 | 68.3 | 20.0 |
| S08 | 41 | M | 2.4.5 | Kb | 8 | 10 | 145 | 143 | 75.6 | 15.9 |
| S09 | 52 | M | 2.4.7 | RTBox* | 8 | 8 | 444 | 446 | 83.9 | 15.6 |
| S10 | 32 | M | 2.4.8 | RTBox | 1 | 5 | 286 | 282 | 81.2 | 17.4 |

**Figure 1:**
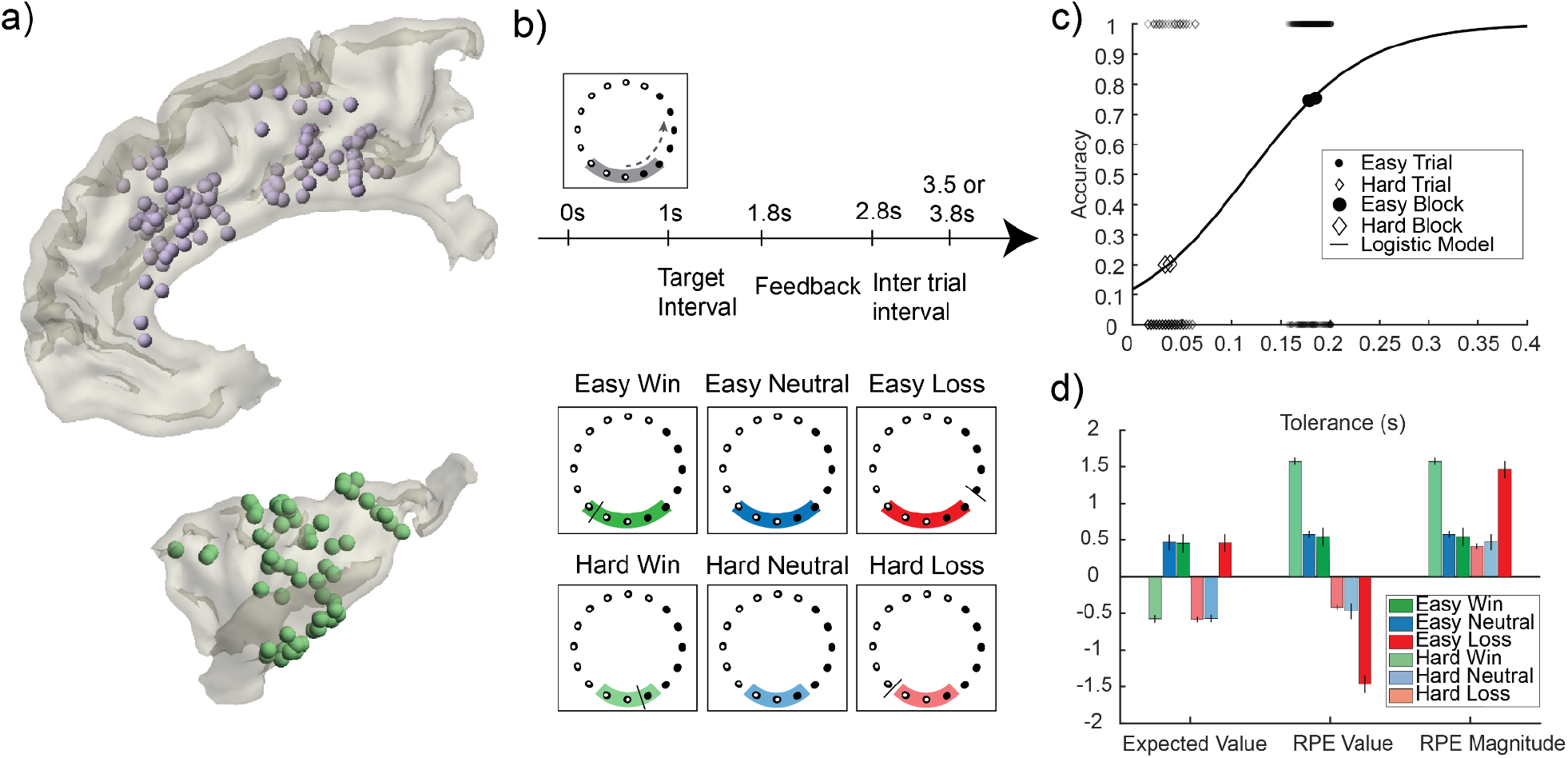
Task design, behavioral modeling, and iEEG recording sites. (a) Reconstruction of iEEG recording sites in dMPFC (top) and INS (bottom) across all participants plotted on a standardized group brain after mirroring all channels to the right hemisphere for dMPFC and left hemisphere for INS. Participants pressed a button to estimate the time when lights finished moving around a circle. The gray target zone cue displayed error tolerance around the 1 s target interval. Audiovisual feedback is indicated by the tolerance cue turning green for wins and red for losses. A black tick mark displayed RT feedback. For 4 patients, blue neutral feedback was given with no RT marker on 12% of randomly selected trials. (c) Tolerance and outcome data for an example participant. Larger markers show block level accuracy; smaller markers show binary single trial outcomes. Model fit using logistic regression provides single trial estimates of win probability, which is converted to expected value. (d) Predictions for RL model predictors. Error bars indicate standard deviation between participants.

### Behavioral modeling

In order to quantify valenced and non-valenced RPE features, we used computational modeling of individual patient behavior to derive single-trial estimates of expected value, RPE value, and RPE magnitude. For each patient, we used logistic regression to predict binary win/loss outcomes across the entire session using error tolerance (Fig. 1c). This model yields patient-specific win probabilities for a given tolerance, which was linearly scaled to the reward function (1, 0, or -1 for winning, neutral, or losing outcomes) to quantify expected value for every trial. Single-trial RPE values were computed by subtracting the expected value from the outcome value, and RPE magnitudes were defined as the absolute value of RPEs. Notably, different reward expectations across easy and hard conditions shift the RPE valence of neutral outcomes to negative in easy blocks and positive in hard blocks (see model predictions in Fig. 1d).

### Positive and negative RPE are encoded in a separate, valence-specific manner

To determine whether neurons encode RPE value, RPE magnitude, or a distribution of positive and negative RPEs, we assessed how well different sets of RL variables predicted the neural data. Towards this aim, we extracted and normalized high frequency band activity (HFA) power from 70-150 Hz at each electrode in dMPFC and INS as a proxy for local population activity (Fig. 2a)^48,51^. Single-trial HFA power was averaged in 50 ms windows sliding by 25 ms from 0 to 600 ms after feedback onset, and these averaged HFA power values were predicted by the different RL variables using linear mixed-effects models across channels and subjects per region and window.

**Figure 2:**
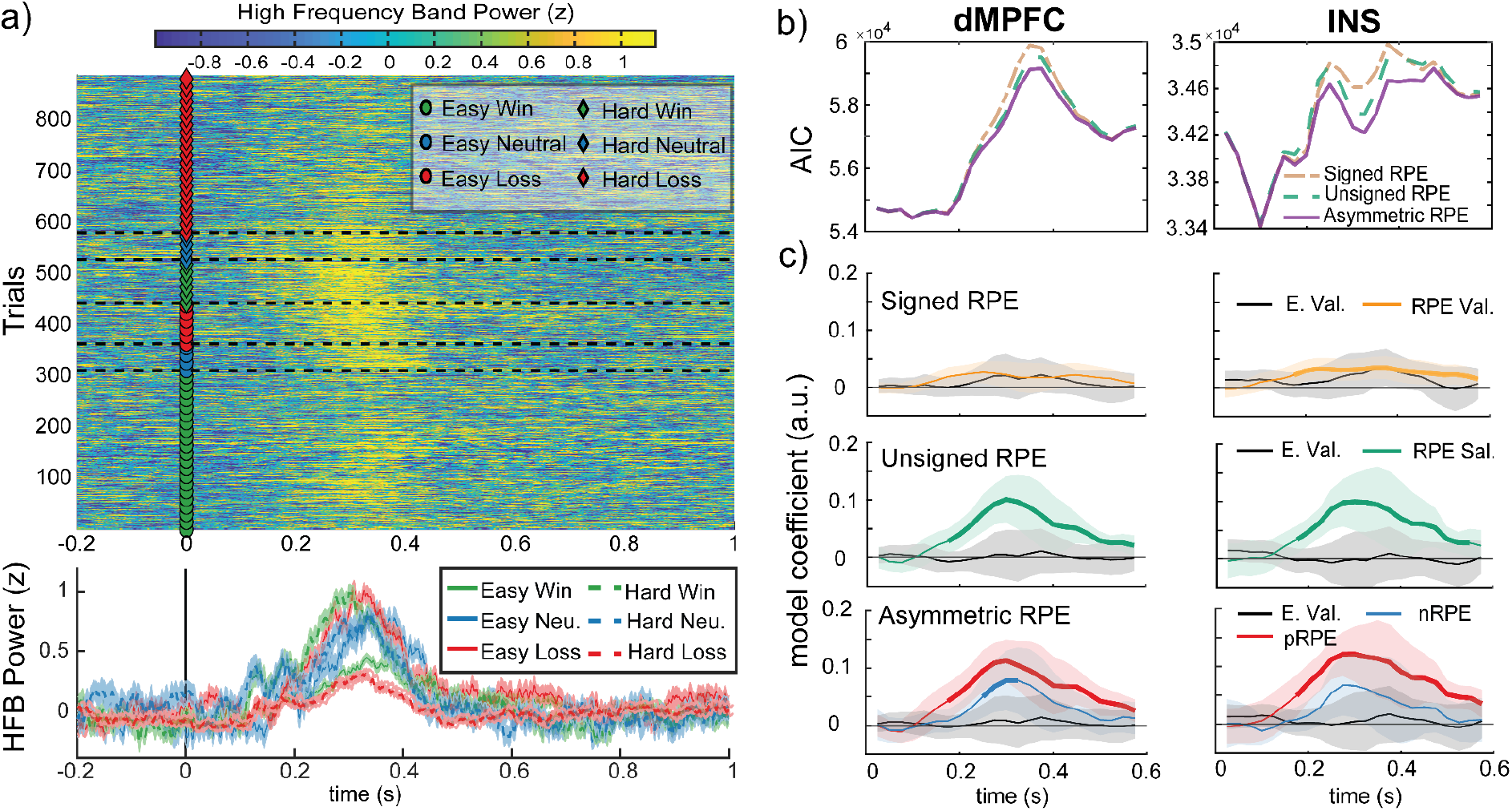
Positive and negative RPE are encoded in a separate, valence-specific manner. (a) Top: Single-trial HFA power at an example channel in the INS is plotted time-locked to feedback (markers at feedback indicate condition). Bottom: Condition averaged HFA power (error bars represent standard error of the mean). (b) Model performance comparison between the three sets of RL variables using Akaike Information Criterion (AIC) for each HFA power time window. Lower values indicate better performance. Region-level, fixed-effects coefficients from linear mixed-effects models predicting single-trial HFA power with three different sets of RL variables: Signed RPE (expected value + RPE value), unsigned RPE (expected value + RPE salience), and asymmetric RPE (expected value + positive RPE + negative RPE). HFA power was averaged in 50 ms sliding windows (step size of 25 ms). Significant model coefficients (q_FDR_ < 0.05) are plotted in bold. Error bars correspond to 95% confidence intervals.

The resulting fixed-effects model coefficients for each window provide a time series depicting the evolution of the different predictors for a given region.

For both regions, the asymmetric RPE model predicted HFA power best, followed by RPE salience and then RPE value (Fig. 2b). Model coefficients (Fig. 2c) indicate RPE value was significantly above zero only in INS, while RPE salience was significantly above zero in both regions. This suggests HFA power increases with larger RPE values and magnitudes. However, the asymmetric model coefficients indicated that positive RPE and negative RPE are encoded differently. Positive RPE is significantly associated with an increase in HFA power in both regions, peaking around 275 ms in INS and 300 ms in dMPFC after feedback onset. In contrast, the negative RPE effect, although qualitatively similar, was weaker in both regions and non-significant in INS. The fact that the asymmetric model performs best indicates that RPE value and RPE magnitude alone cannot explain HFA activity and that neuronal populations exhibit asymmetric coding of negative and positive RPEs.

### Diverse responsiveness of neuronal populations to negative and positive RPE

To understand how different RPE features are coded by neuronal populations in each region, we classified channels with significant responses to positive and/or negative RPEs into four categories (Fig. 3a)^32^: 1) **positive RPE** (increasing/decreasing HFA power with positive RPE magnitude and no significant response to negative RPEs); 2) **negative RPE** (increasing/decreasing HFA power with negative RPE magnitude and no significant response to positive RPEs); 3) **signed RPE** (increasing HFA with positive RPE and decreasing HFA power with negative RPE, or vice versa); and 4) **unsigned RPE** (increasing/decreasing HFA power with both positive and negative RPE magnitude). Note that, for each category, RPE can be encoded with both decreases and increases in HFA, which poses a challenge to both classical RL and dRL. In both these theories, positive RPEs are represented in activity increases, whereas negative RPEs are represented in decreases (henceforth called “regular coding”). This means that classical and dRL account for only three of the eight possible coding strategies shown in Fig. 3a. Other strategies, such as unsigned RPE coding and cases in which neurons decrease their firing to positive RPE and increase it to negative RPE (henceforth called “inverted coding”) are not incorporated in these theories. In the following, we evaluate the extent to which different coding strategies arising from asymmetric scaling are present in the salience network.

**Figure 3:**
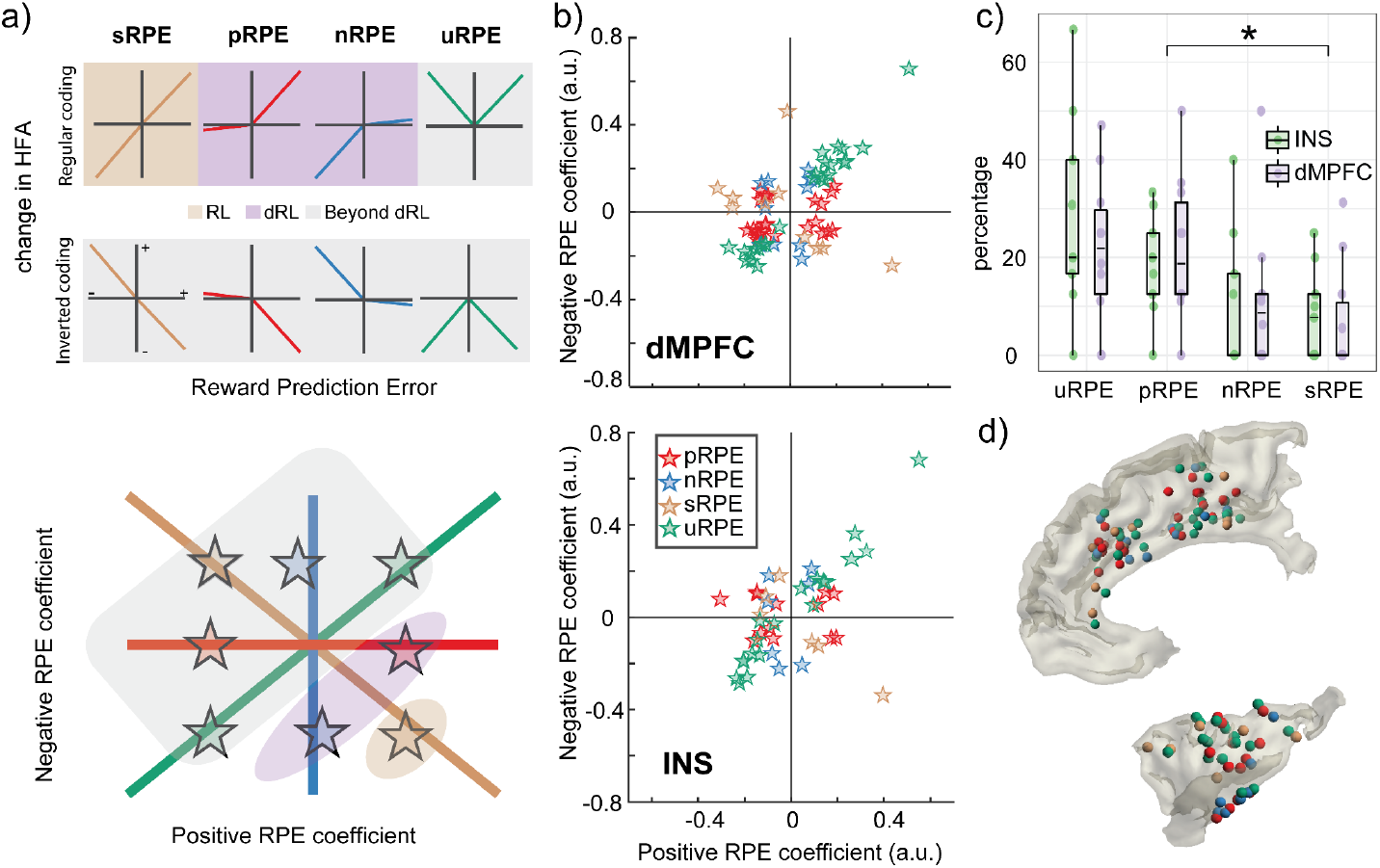
Diverse responsiveness of neuronal populations to negative and positive RPE. (a) Schematic of the different responsiveness profiles of local neuronal populations (top), classified into four categories: Positive (p) RPE (red), negative (n) RPE (blue), unsigned (u) RPE (green), and signed (s) RPE (gold) (see main text for details). For each category, the coding strategy could be regular (i.e., increasing/decreasing HFA power with increasing positive/negative RPE magnitude) or inverted (i.e., decreasing/increasing HFA power with increasing positive/negative RPE magnitude). uRPE populations are labeled by whether they increase or decrease activity regardless of valence. Note that classical RL (gold shade) and distributional RL (purple shade) only account for regular sRPE, pRPE, and nRPE responsiveness. In contrast, unsigned RPEs (i.e. RPE salience) and inverted coding strategies (gray shade) emerge from asymmetric scaling but are not incorporated in current theories. Populations responding to the four categories (colored stars) can be projected on a positive vs. negative RPE plane (bottom). Note how pRPE and nRPE units spread along the x and y axes, while uRPE and sRPE units spread along the diagonal and off-diagonal. (b) Positive and negative RPE peak coefficients for responsive channels belonging to each of the four categories, projected on the two-dimensional RPE plane. (c) Proportion of channels per participant and ROI falling within each category. Box plots depict median and interquartile range. (d) Anatomical location of responsive channels colored per category (Top: dMPFC, bottom: INS). All channels were mirrored to the right hemisphere for dMPFC and left hemisphere for INS.

In both regions, the most frequent response profile encoded unsigned RPE (i.e. RPE salience), with a median of 20.00% (IQR = 15.62-42.5) of channels in INS and 21.88% (IQR = 11.11-31.25) of channels in dMPFC (Fig. 3b). A significant proportion of channels encoded positive RPE only, with a median of 20.00% (IQR = 11.88-26.44) in INS and 18.75% (IQR = 12.5-33.33) in dMPFC. A lower proportion encoded negative RPE only (MDN = 0.00%, IQR = 0.00-18.5 in INS and MDN = 8.68%, IQR = 0.00-12.5 in dMPFC), and a minority of channels encoded signed RPE (i.e., RPE value; MDN = 7.69%, IQR = 0.00-14.37 in INS and MDN = 0.00%, IQR = 0.00-12.5 in dMPFC). When pooling all subjects together, there were 24 (23%) purely positive RPE channels, 10 (9%) purely negative RPE channels, 10 (9%) signed RPE channels and 26 (25%) unsigned RPE channels among the 106 sites in dMPFC. Similarly, there were 13 (20%) purely positive RPE channels, 7 (11%) purely negative RPE channels, 6 (9%) signed RPE channels and 19 (30%) unsigned RPE channels among the 64 sites in INS. We did not find significant differences in category proportions between regions (all *q*_*FDR*_ > .49), suggesting similar coding schemes in INS and dMPFC (Fig. 3c). However, there were significant differences between categories in the proportion of responsive channels when averaged across regions, *χ*^2^(3) = 10.5, p = .01. Post-hoc, pairwise comparisons revealed lower proportions for signed RPE compared to positive RPE (*q*_*FDR*_ = .047), and compared to unsigned RPE only before FDR correction (p = .042). No other significant differences were found between categories (all *q*_*FDR*_ > .30). All categories were spatially interleaved, indicating mixed coding of RPE features across the cortical surface of both regions (Fig. 3d).

Next, we evaluated the extent to which different channels exhibited inverted coding strategies as defined above. We found that 23/37 (62.2%) of positive RPE channels and 9/16 (56.3%) of signed RPE channels decreased their activity with increasing positive RPE magnitude, while 10/17 (58.8%) of negative RPE channels increased their activity with increasing negative RPE magnitude. Similarly, 22/45 (48.9%) of unsigned RPE channels decreased activity with both positive and negative RPE magnitude. This indicates that key variables such as RPE salience and value can be represented by populations of neurons that separately code for negative and positive RPE using both increases and decreases in activity, resulting in inverted coding schemes with respect to classical RL and dRL.

### RPE variables predominantly modulate directed connectivity from INS to dMPFC

Given previous reports indicating that INS might lead information transfer in the network, we next asked how different RPE variables were communicated between regions by estimating directed functional connectivity between INS and dMPFC. Using cross-correlation, we calculated, for each participant, how well activity in each channel of one region predicted the activity of each channel in the other region, at different time lags. We found that, at the region level, positive and negative RPE magnitude increased correlation between INS and dMPFC channels with a peak lag of 75 ms, meaning that INS activity predicted dMPFC activity best at a 75 ms delay (Fig. 4a).

**Figure 4.**
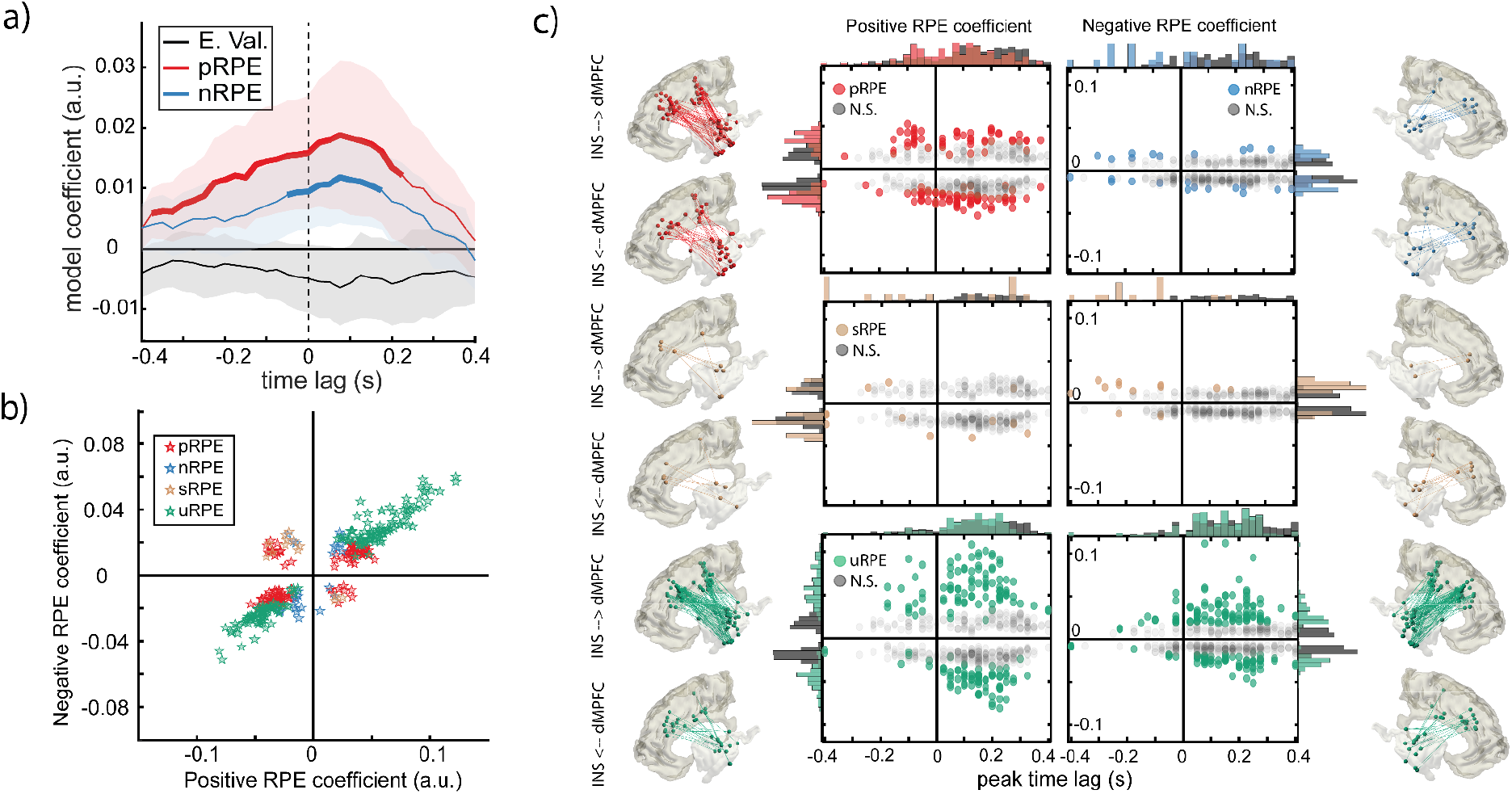
RPE features predominantly modulate directed connectivity from INS to dMPFC. a) Region-level, fixed-effects coefficients depicting the effect of expected value (E. Val.), positive (p) RPE, and negative (n) RPE on the correlation between INS and dMPFC activity at different time lags. Positive lags indicate INS activity precedes dMPFC activity, whereas negative lags indicate dMPFC activity precedes INS activity. Significant model coefficients (q_FDR_ < 0.05) are plotted in bold. Error bars correspond to 95% confidence intervals. b) Negative RPE and positive RPE peak coefficients grouped by category: Positive (p) RPE (red), negative (n) RPE (blue), unsigned (u) RPE (green) and signed (s) RPE (gold). c) Peak coefficients and the respective peak time lags, grouped by category. Significantly responsive channel pairs are displayed in color. Marginal distributions of peak time lags and coefficients are plotted on x and y axes, respectively. The anatomical location of significant channel pairs is shown for each category, separately for the two directions of communication. Channel positions have been projected to the right/left hemisphere for dMPFC and the left/right hemisphere for INS in the case of positive/negative RPE coefficients.

To investigate communication of RPE variables, we classified between-region channel pairs into the same four categories used for HFA analyses. In this case, channel pairs that significantly decreased or increased their correlation as a function of negative and/or positive RPE were classified according to their peak coefficient value. We found significant differences between categories in the proportions of channel pairs *χ*^2^(3) = 9.28, p = .03 (Fig. 4b), with a majority responding to unsigned RPE (MDN = 17.19%, IQR = 0.00 - 23.77) followed by purely positive RPE (MDN = 12.00%, IQR = 7.12 - 20.05). Fewer pairs responded to purely negative RPE (MDN = 2.08%, IQR = 0.00 - 7.78) and a minority responded to signed RPE (MDN = 0.00%, IQR = 0.00 - 1.18). Pairwise contrasts between categories revealed significant differences between pRPE and sRPE after FDR correction (*q*_*FDR*_ = .047), and between sRPE and both uRPE (p = .047) and nRPE (p = .031) before correction. This pattern of results is similar to that found in HFA analyses.

To investigate whether the direction of communication was different across RPE features, we next tested for differences in peak lags between RPE categories. We found that lags were predominantly positive, with uRPE having the longest median peak lag (MDN = 150 ms, IQR = 50 - 200) for positive RPE coefficients, followed by pRPE (MDN = 80 ms, IQR = -50 - 180) and then sRPE (MDN = -30 ms, IQR = -290 - 280). For negative RPE coefficients, uRPE had again the longest median peak lag (MDN = 150 ms, IQR = 80 - 230 ms) followed by nRPE (MDN = 30 ms, IQR = -180 - 230) and then sRPE (MDN = -180 ms, IQR = -240 - -80). This suggests that information predominantly flowed from INS to dMPFC (Figure 4c). However, we found significant differences in peak lags among categories for negative (*χ*^2^(3) = 31.48, p < .001) but not positive (*χ*^2^(3) = 5.33, p = .15) RPE coefficients. These differences were mainly driven by sRPE (*q*_*FDR*_ = .004) and nRPE (*q*_*FDR*_ = .003) lags being significantly more negative than uRPE lags. These results suggest potential bidirectional communication for these categories such that sRPE and nRPE may have also been communicated from dMPFC to INS.

In contrast to region-level results, where lagged correlations increased with positive and negative RPE magnitude, there was a significant portion of individual pairs whose correlation showed an inverted coding scheme as defined above. We observed 70/121 (58%) of pRPE pairs and 10/13 (77%) of sRPE pairs decreased their correlation with increasing positive RPE magnitude, while 10/24 (42%) of nRPE pairs increased their correlation with increasing negative RPE magnitude. Similarly, 72/167 (43%) of uRPE pairs decreased their correlation with both positive and negative RPE magnitude. As in single channel responsiveness, this finding emphasizes the utility of decreases in HFA as a means to encode information. Finally, channel pairs involved in RPE communication between INS and dMPFC were also spatially interleaved, which agrees with the aforementioned mixed coding scheme for RPEs in neuronal populations (Fig 4c).

## DISCUSSION

The valence and salience of RPEs are critical components of reinforcement learning and cognitive control. However, it is unclear how dMPFC and INS networks represent these variables to facilitate behavioral adaptation. Using HFA power as a proxy for local population activity, we show that a model utilizing asymmetric positive and negative RPE coding explained feedback-related activity in dMPFC and INS better than models including only RPE value or RPE salience. While positive RPE signals were robustly encoded in both regions, negative RPE signals were less prominent and only significant in dMPFC, suggesting a bias towards positive reward outcomes in these areas. Moreover, neuronal populations at individual channel sites exhibited distinct responsiveness profiles, allowing flexible encoding of RPE value and salience with both increases and decreases in activity. A plurality of channels responded to RPE salience (25% in dMPFC, 30% in INS) and purely positive RPEs (23% in dMPFC, 20% in INS). A lower proportion of sites encoded negative RPEs (9% in dMPFC, 11% in INS), and few encoded signed RPE (9% in dMPFC, 9% in INS). This indicates that non-linear, heterogeneous representations of reward information are the dominant coding scheme in dMPFC and INS. Finally, directed connectivity measures indicated channel pairs were primarily modulated by positive RPEs and RPE salience, and that communication for these variables flows predominantly from INS to dMPFC. Collectively, these results demonstrate that neuronal populations respond differently to positive and negative RPEs, enabling RPE coding diversity in human dMPFC and INS. Below, we discuss how these findings inform theoretical debates in reward learning and cognitive control and their conceptual and methodological implications for principles of neural coding.

Our findings that a model with valence-specific RPEs better explains dMPFC and INS activity has several implications for theories of neural coding and function in the salience network. First, they align with and expand upon recent advances in computational and systems neuroscience by showing that asymmetric scaling principles underlying dRL models better explain heterogeneous responses to reward and punishment. To date, dRL has mainly been applied to explain the diverse response profiles of single units in subcortical dopaminergic circuits of rodents^28,29^. Our findings provide novel evidence that neuronal populations in the human cortex exhibit asymmetric coding, one of the core principles behind dRL.

Our analyses also revealed that reward and salience information were represented with both increases and decreases of HFA activity and connectivity. Distinguishing these opponent coding schemes is important to avoid confounding interpretations of signed RPE value, which may have contributed to conflicting findings in previous studies^46^. However, current dRL models operationalize positive/negative RPEs as increases/decreases in neuronal activity, respectively^28^. This implies that dRL does not capture RPE salience effects or inverted coding schemes, which were prominent in our human cortical data and have also been reported in dopaminergic midbrain neurons^23,27,52^. Our findings highlight the need to devise biologically consistent implementations of asymmetric scaling and reward coding within the dRL framework.

Importantly, we leveraged the flexible coding scheme allowed by asymmetric scaling to categorize combinations of positive and negative coefficients corresponding to the key RPE value and salience variables underlying central theories of dMPFC function. We found that a plurality of individual sites within and connectivity pairs between dMPFC and INS responded to the salience of RPEs, including many of the strongest responses. This observation supports the proposed central role of dMPFC and INS network in coding salience to enable adjustments of cognitive control^5–7,9,17,19,53^. In contrast to signed RPE theories of dMPFC function^10–13^, valenced RPE information was rarely encoded in either neural activity or connectivity responses in a manner consistent with signed RPE value. Notably, our findings are compatible with accounts suggesting dMPFC integrates the positive, negative, and salience RPE variables required to update cognitive control, since separate representations of this information can still be flexibly read out by downstream regions to support adjustments in approach, avoidance, or motivation. However, our data suggest that most neural populations in dMPFC and INS do not represent these variables together as a combined “common currency” value signal with symmetric but opposite coding of positive and negative RPEs. This finding is consistent with recent proposals that neural representations of value are better understood as related to attention, action plans, vigor, or other choice-related variables^54–56^.

Another important result from our single-channel analyses is that diverse populations coding for different RPE variables are spatially interleaved within each region. This observation revealed a more nuanced picture than region-level analyses, which can obscure local heterogeneity within regions. Reconciling population- and region-level results may explain seemingly contradictory evidence supporting different theories of dMPFC function, particularly between single unit studies in systems neuroscience and experiments using functional magnetic resonance imaging or EEG in cognitive neuroscience, which typically average over intermingled populations. Indeed, our population-level results align with nonhuman primate studies that identified single units sensitive to valence-specific and unsigned salience RPEs within dMPFC^11,31,32^, suggesting these circuits are anatomically nonseparable^33^. In contrast, a recent human iEEG study reported an anatomical dissociation between positive and negative RPE processing^36^. However, this study reported region-level effects, leaving the diversity in local population coding of RPE salience and valence unexplored. This emphasizes the need to disentangle spatially intermingled circuits performing different computations within a given region, which is characteristic of previous human iEEG findings in language and attention^57,58^. Furthermore, we found that the proportions of channel sites coding RPE salience, RPE value, and positive and negative RPEs were equivalent in dMPFC and INS. This result supports the view that neural computations underlying reward learning, value-based decision making, and cognitive control unfold in parallel across distributed circuits^54,59–61^.

Our results also indicate that population activity within and connectivity between dMPFC and INS have stronger representations for positive than negative RPEs. This is particularly striking given that our task employs an implicit win-stay/lose-switch strategy, indicating that this bias towards positive RPE coding was observed despite the fact that no adjustments to cognitive control or behavioral responses are required after positive feedback in our task. This finding fits with evidence from nonhuman primate single unit studies^11,32,35^ and some human iEEG results^62^. However, a variety of conflicting evidence from other prior studies argues dMPFC and INS show a bias for processing negative valence^16,36,40,63^. In particular, Gueguen et al. 2021 report a bias for negative RPEs in HFA responses in anterior INS^36^. One methodological source of this discrepancy could be that not all prior studies controlled for salience, as many of the individual and pairs of channels that responded to negative RPEs in our analyses were revealed to code for salience once we accounted for their response to positive RPEs. Alternatively, differences in analyses (e.g., region-versus population-level) or task design, such as the use of positive versus negative punishment (i.e., delivering aversive stimuli versus omitting positive rewards) or interactions between effort and reward^64^, could contribute to these different results.

Another potential factor influencing the proportion of positive, negative, and salience responses is where our specific recording sites are located relative to functional gradients within dMPFC and INS. For example, the strong representations of salience in our results is likely influenced by the majority of our recording sites falling in mid-cingulate and insular cortices overlapping with the salience network, which is associated with control and performance monitoring^9,53,65,66^. In contrast, single units recorded from more anterior regions of MPFC in non-human primates show reduced salience coding and mostly responded to positive and negative RPEs^32^. This difference in the relative strength of signed and unsigned RPE coding is potentially because anterior cingulate cortex is a distinct subregion of MPFC more closely linked to limbic circuits involved in learning, comparing, and choosing values than action control^20,66–69^. Similarly, our results showed some negative RPE coding in the INS that aligns with previous studies reporting a bias towards negative RPEs in the anterior portion of this region^4,36,63,70,71^. However, our spatial sampling of the INS—which was determined solely based on clinical needs of the patient—revealed a bias towards positive RPE representations in mid- and posterior INS. Interestingly, this potential shift in sensitivity from negative to positive bias across the anterior-posterior axis fits with observations from rodent research of a hedonic “hot spot” in the INS where stimulation induces “liking”, which is found posterior to a hedonic “cold spot” in more anterior INS^72,73^. Overall, our converging results from both individual channels and between-region connectivity indicate that dMPFC and INS are predominantly modulated by positive RPEs and RPE salience.

Lastly, the results of our directed connectivity analyses revealed INS-to-dMPFC communication for positive RPEs and RPE salience processing, which provides direct evidence for hypotheses that the INS plays a leading role in the salience network^42,74,75^. Our findings build upon two recent human iEEG studies showing INS-to-dMPFC connectivity for salience^43,44^. However, these studies used tasks that did not dissociate the valence and salience of feedback. Here, we demonstrate that INS-to-dMPFC directed connectivity predominantly conveys both salience and positive RPE information. Thus, in addition to facilitating salience processing between these two control regions, INS-to-dMPFC communication of positive RPEs may reflect integration of affective information from ventral reward systems including the INS into action processing in dorsal control systems including mid-cingulate cortex^76,77^. Unfortunately, too few channel pairs were significantly modulated to draw firm conclusions about the directionality of negative RPE and RPE value communication. These results confirm and expand the role for INS as a general source for multiple RPE variables processed in dMPFC, thereby emphasizing the need for the field to shift from an excessive focus on dMPFC towards including the INS in empirical research and theory building.

In conclusion, our results demonstrate that incorporating asymmetric scaling principles inspired by dRL can capture positive, negative, and salience RPE coding in human dMPFC and INS. Moreover, individual channel analysis strategies similar to those used in non-human systems neuroscience revealed that these populations are interleaved in anatomically overlapping circuits within dMPFC and INS. Importantly, we found that accounting for valence-specific RPE coding using both increases and decreases in activity established that few sites or channel pairs were modulated by signed RPE, arguing against hypotheses that these regions integrate reward and punishment into a common value signal. Instead, our results support a combination of valence-specific and unsigned salience theories of dMPFC and INS function. Finally, our directed connectivity results emphasize the leading role of the INS in both positive and unsigned RPE processing. Overall, these findings bridge region-level analyses common in human neuroscience with population-level analyses in animal models to support and expand dRL principles and reconcile long standing theoretical debates regarding neural coding of RPEs in dMPFC and INS.

## ACKNOWLEDGEMENTS

We thank the participants for their invaluable efforts and I. Griffith for help piloting the paradigm. This work was supported by NINDS R37NS21135 (RTK), CONTE Center PO MH109429 (RTK), Brain Initiative U19NS107609-03 and U01NS108916 (RTK, JJL), NSF GRFP (CWH), and the Independent Research Fund, Denmark (DQM).

## CITATION DIVERSITY STATEMENT

Recent work in several fields of science has identified a bias in citation practices such that papers from women and other minority scholars are under-cited relative to the number of such papers in the field. Here we sought to proactively consider choosing references that reflect the diversity of the field in thought, form of contribution, gender, race, ethnicity, and other factors. First, we obtained the predicted gender of the first and last author of each reference by using databases that store the probability of a first name being carried by a woman. By this measure and excluding self-citations to the first and last authors of our current paper, our references contain 6.33% woman(first)/woman(last), 14.85% man/woman, 20.25% woman/man, and 58.56% man/man. This method is limited in that a) names, pronouns, and social media profiles used to construct the databases may not, in every case, be indicative of gender identity and b) it cannot account for intersex, non-binary, or transgender people. Second, we obtained predicted racial/ethnic category of the first and last author of each reference by databases that store the probability of a first and last name being carried by an author of color. By this measure (and excluding self-citations), our references contain 13.51% author of color (first)/author of color(last), 12.84% white author/author of color, 16.03% author of color/white author, and 57.61% white author/white author. This method is limited in that a) names and Florida Voter Data to make the predictions may not be indicative of racial/ethnic identity, and b) it cannot account for Indigenous and mixed-race authors, or those who may face differential biases due to the ambiguous racialization or ethnicization of their names. We look forward to future work that could help us to better understand how to support equitable practices in science.

## METHODS

### Participants

Data was collected from ten patients undergoing neurosurgical treatment for medically refractory epilepsy (mean ± SD [range]: 35.2 ± 13.1 [21-57] years old; 1 woman; see Table 1 for patient demographics and electrode coverage). Patients were implanted with stereotactic (SEEG) or subdural grid or strip (ECoG) electrodes, and electrode placement and medical decisions were determined solely by the clinical needs of the patient. Patients were observed in the hospital for approximately a week, and those willing to participate performed the Target Time behavioral task during breaks in their clinical treatment. Informed consent was obtained according to experimental protocols approved by the University of California, Berkeley, University of California, Irvine, and California Pacific Medical Center Committees on Human Research. Patients had normal IQ (>85) and spoke fluent English.

### Target Time Behavioral Task

The Target Time interval timing task was written in PsychoPy^78^ (v1.85.3) and consisted of four blocks (two easy and two hard) of 75 trials each (see Fig 1A for task schematic). Two patients completed the task twice, and one patient completed the task three times. The order of block difficulty was fixed as either two easy followed by two hard or alternating from easy to hard (Table 2). Following central fixation and a randomly chosen inter-trial interval ranging from 0.2 to 1.2 s (see Table 2), trials began with presentation of a visual motion cue at a constant speed to arrive at a target at the one-second temporal interval. Participants estimated the interval via button press using the space bar on a keyboard or an RTBox (v5/6) response device ^79^. In the first version of the task (n = 6), the motion cue was upwards in a straight line towards a bullseye target, and in a second version (n = 4), the motion cue was counter-clockwise starting and ending at the bottom of a ring of dots on which a gray target zone was centered. The size of the bullseye and the width of a gray target zone indicated the tolerance for successful responses. Veridical win/loss feedback was presented from 1.8 s to either 2.6 or 2.8 s (Table 2) and composed of (1) the tolerance cue turning green/red, (2) cash register/descending tones auditory cues, and (3) a black tick mark denoting the response time (RT) on the ring. Participants received ±100 points for wins/losses. Tolerance was bounded at ± 15-200 or 15-400 ms (Table 2), and separate staircase algorithms for easy and hard blocks adjusted tolerance by -3/+12 and -12/+3 ms following wins/losses, respectively. Participants learned the interval in five initial training trials in which visual motion completed the full linear track or circle. For all subsequent trials, dot motion halted after 400 ms to prevent visuo-motor integration, forcing participants to rely on external feedback. Training concluded with 15 easy and 15 hard trials to initialize both staircase algorithms to individual performance levels. For the second task version, main task blocks introduced neutral outcomes on a random 12% of trials that consisted of blue target zone feedback, a novel oddball auditory stimulus, no RT marker, and no score change.

**Table 2:**
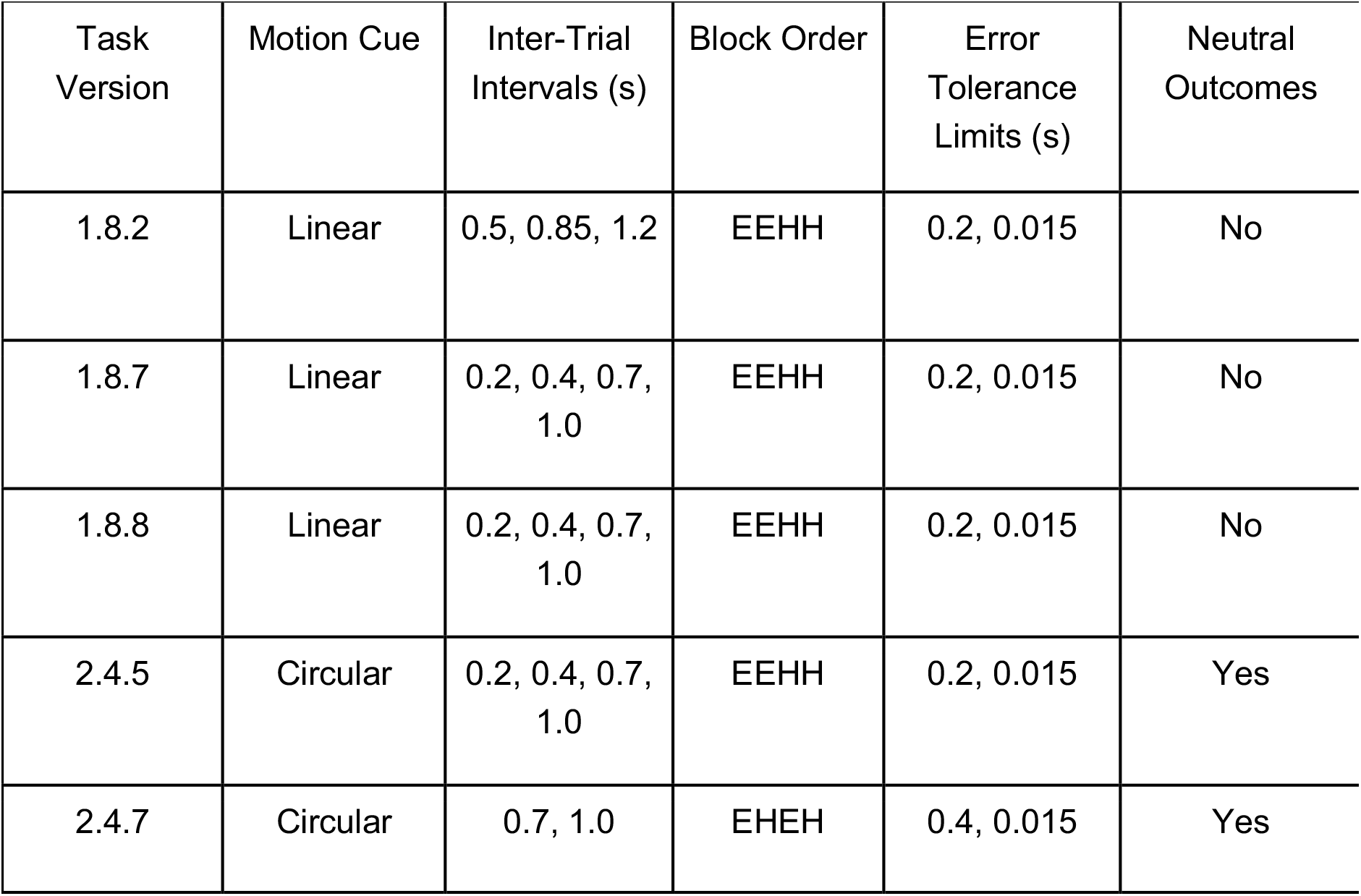
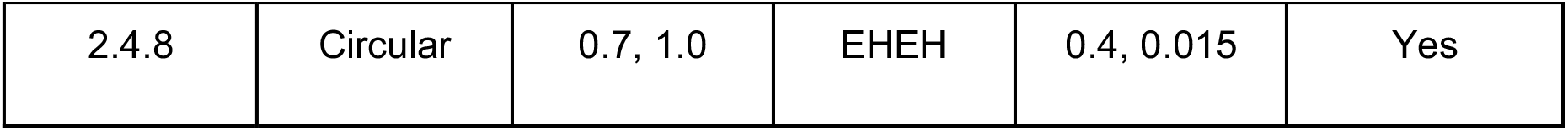
Target Time paradigm parameters. For Block Order, E refers to easy blocks and H refers to hard blocks.

### Behavioral modeling

The relationship between the tolerance around the target interval and expected value was fit to individual participant behavior using logistic regression. Specifically, tolerance was used to predict binary win/loss outcomes across trials using the MATLAB function *glmfit* with a binomial distribution and logit linking function. Trials with neutral outcomes were excluded because they were delivered randomly and thus not reflective of performance. The probability of winning (*p*_*win*_) for each participant was computed as:

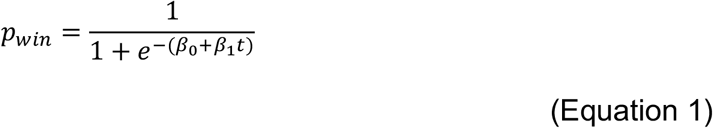

where *β*_*0*_ is the intercept and *β*_1_ is the slope from the logistic regression, and t is the tolerance on a given trial. Expected value was derived by linearly scaling the probability of winning to the reward function ranging from -1 to 1. RPE value was then computed by subtracting expected value from the actual reward value, and RPE magnitude was computed as the absolute value of RPE value. See Figure 1c for model predictions by condition.

### iEEG data collection, localization, and preprocessing

The data were recorded at either the University of California Irvine Medical Center (n = 9), USA or California Pacific Medical Center (n = 1), USA. Patients at Irvine were implanted with stereo-EEG (SEEG) electrodes with 5 mm spacing, and the patient at CPMC was implanted with strips of electrocorticography (ECoG) electrodes with 1 cm spacing. At both sites, electrophysiology and analog photodiode event channels were recorded using a 256-channel Nihon Kohden Neurofax EEG-1200 recording system and sampled at 500 (n = 3), 1000 (n = 3), or 5000 Hz (n = 4). For five patients, analog photodiode channels and a subset of iEEG channels were recorded in a separate Neuralynx ATLAS recording system at Irvine at 4000 (n = 1) or 8000 Hz (n = 4). For these cases, photodiode events were then aligned to the iEEG data acquired in parallel via the Nihon Kohden clinical amplifier via cross-correlation of shared iEEG channels. Pre-operative T1 MRI and post-implantation CT scans were collected as part of standard clinical care, and recording sites were reconstructed in native patient space by aligning these scans via rigid-body co-registration according to the procedure described in Stolk et al. ^80^. Anatomical locations of electrodes were determined by manual inspection in native patient space under supervision of a neurologist. Electrode positions were then normalized to group space by warping the patient MRI to a standard MNI 152 template brain using volume-based registration in SPM 12 as implemented in Fieldtrip ^80^. Group-level electrode positions are plotted in MNI coordinates relative to the cortical surface of the fsaverage brain template from FreeSurfer ^81^.

Data cleaning, preprocessing, and analyses were conducted using the Fieldtrip toolbox ^82^ and custom Python and MATLAB code. Raw iEEG traces were manually inspected by a neurologist for epileptic spiking and spread, as well as artifacts (e.g., machine noise, signal drift, amplifier saturation, etc.). Data in regions or epochs with epileptiform or artifactual activity were excluded from further analyses. Preprocessing included resampling data to 1000 Hz (for datasets recorded at sampling frequencies > 1000 Hz), bandpass filtered using a Butterworth filter from 0.5-300 Hz, re-referenced (bipolar to adjacent electrodes for SEEG data; common average reference across all channels for ECoG data), and bandstop filtered at 60, 120, 180, 240, and 300 Hz (Butterworth filter with 2 Hz bandwidth) to remove line noise and harmonics. Continuous data were then visually inspected to ensure all epochs with artifacts or spread from epileptic activity were removed. Finally, trials were rejected for task interruptions and behavioral outliers (RTs missing, < 0.5 s, > 1.5 s, or > 3 standard deviations from that patient’s mean), resulting in 274-890 trials per patient (mean ± S.D.: 405.0 ± 210.6).

### High frequency broadband power extraction and modeling

Time series data were filtered to high frequency band activity (HFA) ranges known to correlate with local multi-unit activity ^48,50,51^. Specifically, data were segmented from -0.25 to 1.2 s relative to feedback onset, and multitaper time-frequency transformations with 50 ms windows were used to extract power from sub-bands ranging from 70 to 150 Hz in 10 Hz steps. These HFA power values were then log transformed to account for their log-normal distribution ^83^ in preparation for linear modeling. To normalize these power values against baseline activity, permutation distributions were created for each channel by taking the mean and standard deviation of baseline power values from -0.25 to -0.05 s relative to stimulus onset from 500 iterations of sampling trials with replacement. Feedback-lock power values were then z-scored using the average mean and standard deviation values from those permutation distributions of pre-stimulus baseline power values. This process avoids normalizing HFA power to pre-feedback data which may contain post-response activity and is robust to noisy outlier trials that can skew the baseline data. Finally, sub-bands were averaged together to create a single HFA power time series.

A sliding window approach was then used to average normalized single-trial HFA power values in 50 ms windows stepping by 25 ms from 0 to 0.6 s post-feedback. Mixed-effects models with subject and channel as nested random effects were then used to predict single-trial HFA power data for each time window and brain region. Using AIC as a performance metric, we compared three different RL models, all containing expected value and a unique set of RPE estimates as predictors: The **signed RPE** model included valenced RPE magnitude estimates (i.e. RPE value); the **unsigned RPE** model included absolute RPE magnitude estimates (i.e. salience); and the **asymmetric RPE** model included separate predictors for positive and negative RPE magnitude estimates. Note that the asymmetric RPE model is mathematically equivalent to a model in which both RPE value and salience are introduced as predictors. That is, RPE value and salience emerge as a linear combination of positive and negative RPE. The asymmetric model was added to operationalize our hypothesis and improve interpretability. Furthermore, in previous work we have shown that separating positive and negative RPE magnitude helps to disentangle event-related components that are heavily mixed in scalp EEG data ^47^.

Confidence intervals and two-sided *p*-values for both fixed (i.e., region level) and random (i.e., subject / channel specific) effects coefficients were obtained from the standard error estimates for each time window. *p*-values of region-level fixed effects were corrected for multiple comparisons across time using the false discovery rate (FDR) methods of Benjamini & Hochberg ^84^ for each channel. Corrected *p*-values are referred to as *q*_*FDR*_ throughout the manuscript. *p*-values of channel-specific random effects were left uncorrected, since the regularizing properties of mixed-effects models result in conservative coefficient estimates that protect against false positives and overfitting. Channels were considered to be significantly predicted by a model regressor if any HFA power window had a model coefficient with *p* < 0.05.

### Estimation and inference on channel responsiveness categories

We classified the channels into four categories according to their responsiveness. First, we selected **positive RPE** channels as those significantly predicted by positive RPE estimates only. Similarly, **negative RPE** channels were those significantly predicted by negative RPE only. A third category, **signed RPE**, was composed of channels that responded by significantly increasing their activity with positive RPE, while significantly decreasing activity with negative RPE, or vice versa. Finally, we defined **unsigned RPE** channels as those that either increased or decreased their activity in response to both positive and negative RPE magnitude. Because responsiveness changed over time, in a handful of cases a channel could be classified in both the signed and unsigned RPE categories. In those instances, we classified the channel according to the sign of their peak significant coefficients.

To evaluate differences between regions and channel categories, we calculated the proportions of all channels belonging to each category for each subject and region. One subject was excluded from this analysis as they had no electrodes in INS. At the group level, we used Wilcoxon signed-rank tests to compare channel proportions between regions for each category separately. The resulting *p*-values were FDR corrected across the four between-region tests. After confirming no significant differences between regions for any category, we averaged proportions across regions and tested for differences between categories with a Kruskal-Wallis test followed by post-hoc, FDR-corrected pairwise comparisons with Wilcoxon signed-rank tests.

### Estimation of directed connectivity between INS and dMPFC

We estimated the directed functional connectivity between dMPFC and INS using time-lagged cross-correlation of HFA power time series between all channels in one region and all channels in the other region for each subject. Lags ranged from -400 to 400 ms in 25 ms steps. In our case, positive lags indicate activity in INS precedes activity in dMPFC whereas negative lags indicate dMPFC activity precedes INS activity. Zero lag indicates no delay between regions.

The resulting correlation-coefficient time-lag series were then predicted by the asymmetric RPE model including expected value, positive RPE magnitude and negative RPE magnitude as regressors. For each time lag, a mixed-effects model was estimated including subject and channel pair as nested random effects. *p*-values for (region level) fixed effects and (channel-pair level) random effects were obtained based on standard error estimates. For fixed effects, *p*-values were FDR corrected across time-lags for each predictor separately. For random effects, *p*-values were left uncorrected due to the regularizing properties of mixed-effects models.

For each channel pair and region, we extracted the time lags at which positive and negative RPE magnitude best predicted directed connectivity by finding the peak of the absolute correlation coefficients. We classified channel pairs into the same four categories (pRPE, nRPE, sRPE, uRPE) used for HFA analyses, according to their modulation by negative RPE and / or positive RPE, as indicated above. To evaluate differences in category proportions, we calculated the percentage of all channel pairs belonging to each category for each subject. We tested for differences between categories with a Kruskal-Wallis test followed by post-hoc, FDR-corrected pairwise comparisons with Wilcoxon rank-sum tests. The same statistical procedure was followed to test for differences in peak lags between categories.

